# From Face-to-Face to Online Modality: Implications for Undergraduate Learning While the World is Temporarily Closed in the Age of COVID-19

**DOI:** 10.1101/2020.08.30.274506

**Authors:** Gokhan Hacisalihoglu

## Abstract

The second half of the Spring 2020 semester has been an unprecedented time globally due to the ongoing coronavirus disease 2019 (COVID-19). COVID-19 pandemic has forced more than one billion students out of school, has disrupted the world and led all university courses switched to online instruction social distancing actions taken to limit the spread of the virus. The aim of the present study is to evaluate the pandemic related changes for undergraduate students, to assess their perspectives related to their learning, experiences in two courses, and to discuss the far-reaching potential implications for the upcoming summer and fall semesters. An electronic survey was conducted to gather data on the student perceptions and learning characteristics of this transition from face-to-face (F2F) to online at a medium-sized university in the Southeast in the Spring 2020 semester. Nearly 88% of the participants indicated that the COVID-19 pandemic impacted their education, while 19% indicated that they prefer online over F2F learning. Furthermore, the online modality significantly increased attendance in General Biology I. Our study also showed that the usage of live conferencing and digital applications increased due to the pandemic. The current research fills the gap in the existing literature by providing the first study on the effects of the ongoing COVID-19 pandemic on undergraduate learning and experiences in the most unique dual modality of the Spring 2020 semester.

## Introduction

Viruses are very small pieces of organic material that are only visible under an electron microscope and cause diseases in humans, animals, and plants (Urry et al., 2020). The coronavirus disease 2019 (COVID-19), which is caused by severe acute respiratory syndrome 2 (SARS-CoV-2), was first identified in Wuhan, China on December 8, 2019 and resulted in an unprecedented public health crisis (WHO, 2020). The SARS-CoV-2 virus has spike proteins that attach to host receptors (ACE2, angiotensin converting enzyme 2) via droplets within a 2 m range with an estimated reproduction number of 2.68 (R0; CDC, 2020). As of August 30, 2020, the global number of confirmed cases stands at 25 million while the fatal cases stand at 843,586 (CSSE 08/30/20; Fig. 1). After the COVID-19 pandemic was declared by the World Health Organization (WHO, March 11, 2020), universities responded by temporarily closing down campuses to slow the virus’s spread and flatten the curve. The world has changed due to the aggressive spread of the COVID-19 pandemic and all face-to-face (F2F) university courses shifted to online for the first time in the middle of the Spring 2020 semester.

**Figure 1.**
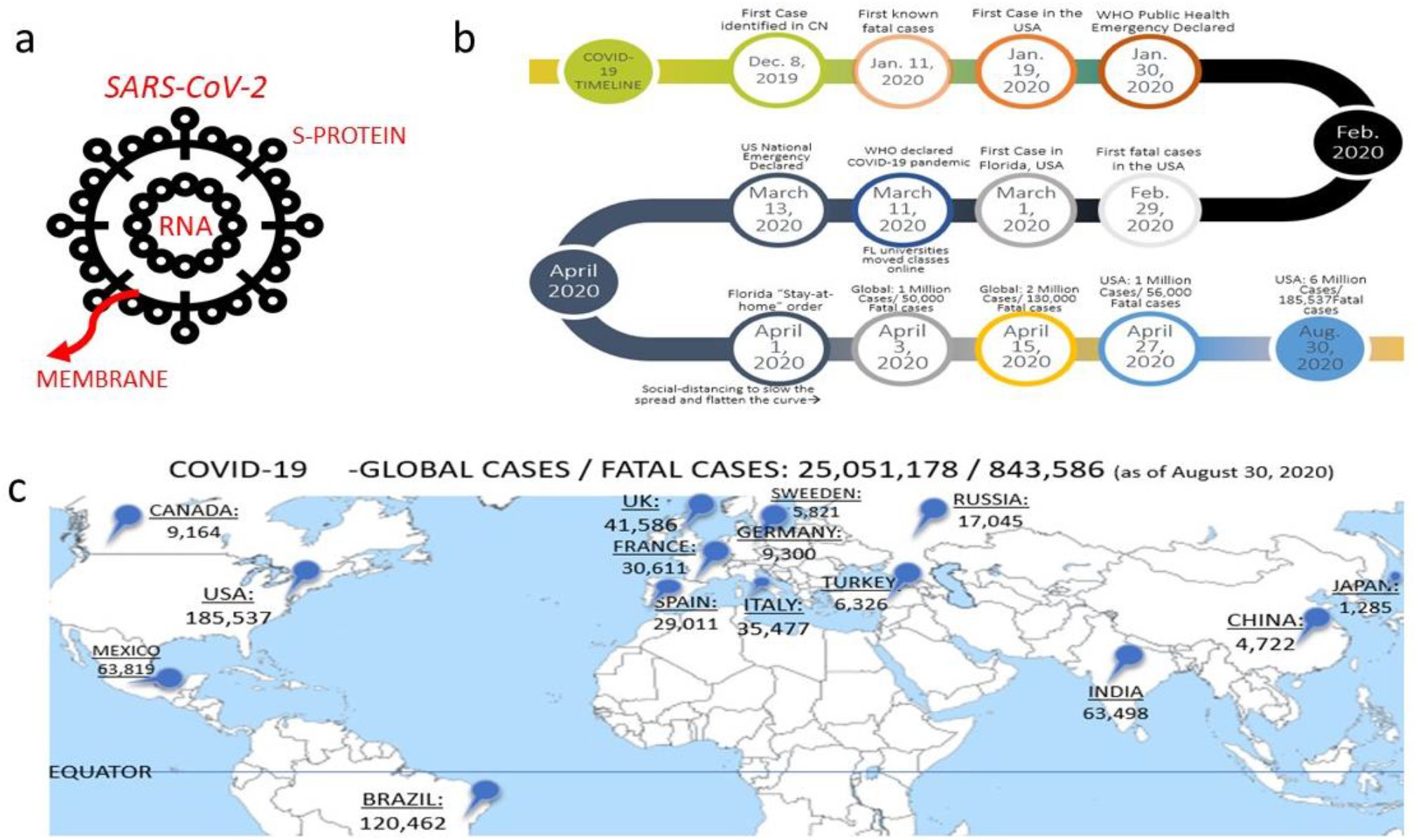
(a) Specifics of the basic structures of coronavirus 2 (SARS-CoV-2); (b) The COVID-19 pandemic timeline; and (c) Global hotspots with severe COVID-19 disease (Data source: CSSE, 2020).

As opposed to F2F, the online modality is characterized by the separation of the instructor and learners in an online environment (Fig. 2; Nilson & Goodson, 2018). Similar to the correspondence distance education programs that started in 1800s, the online modality offers several benefits including learning at anytime and anywhere, a faster completion time, flexible scheduled learning, and lower costs due to the absence of room logistics needs (Fig. 2; Nilson & Goodson, 2018). Moreover, strategies such as live video conference lecturing via Zoom (ZM, San Jose, CA) have been utilized to increase student engagement.

**Figure 2.**
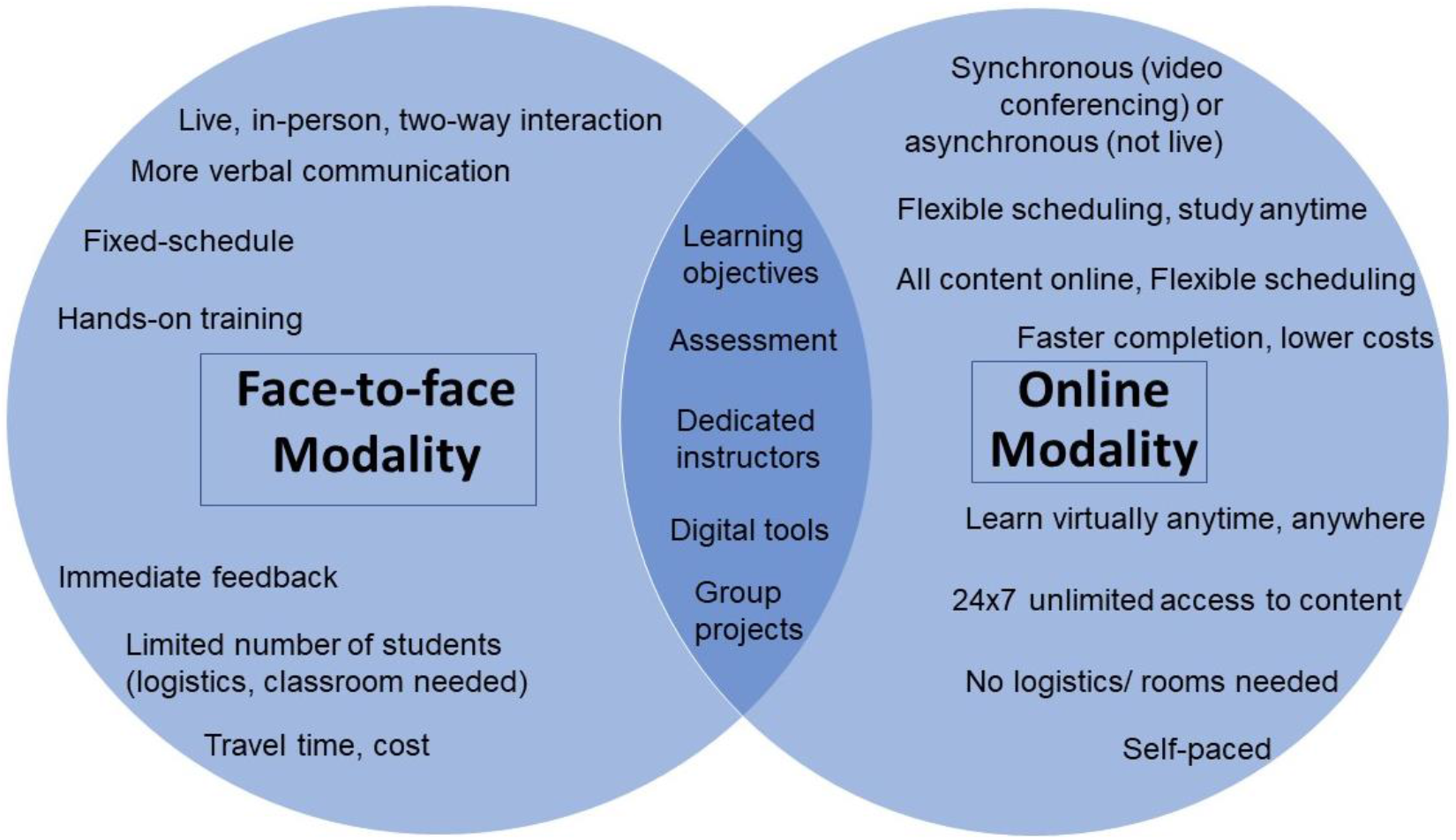
A comparison of the similarities and differences between the face-to-face (F2F) and online learning modalities.

Additionally, F2F and online instruction modalities share many qualities including digital tools, group projects, student learning objectives (SLOs) and assessment (Fig. 2). Nennig et al. (2020) compared student attitudes in online and F2F inorganic chemistry courses. No significant differences were found in performance, grade distribution, and student attitudes. Another study by Mullen (2019) compared F2F with synchronous online learning and found that the online modality was equally as effective. In a medical education study, the researchers showed that both F2F and online formats delivered the same benefits to students in Australia (Ifediora, 2019).

At our university, there is currently a focus on developing more online STEM courses. While previous studies have investigated both modalities, there is no literature on dual modalities in the same semester with one instructor.

Therefore, the objective of this study is to examine the differences in undergraduate student perceptions in F2F versus online STEM biology courses in the unique dual modality of the Spring 2020 semester at Florida A&M University.

## Methods

### Course Structures

The current study was conducted with a cohort of students at a medium-sized university in the Southeast who were enrolled in General Biology I (BSC1010) and Plant Morphology (BOT3303). The Spring 2020 semester was divided into F2F (January to mid-March) and online (mid-March to May) modality sections that were taught by the same instructor. The F2F sections were taught in-person while the online sections were taught using the Zoom (Fig. 3a; Zoom Video Communications [ZM], San Jose, CA) software with video-conferencing and screen sharing.

**Figure 3.**
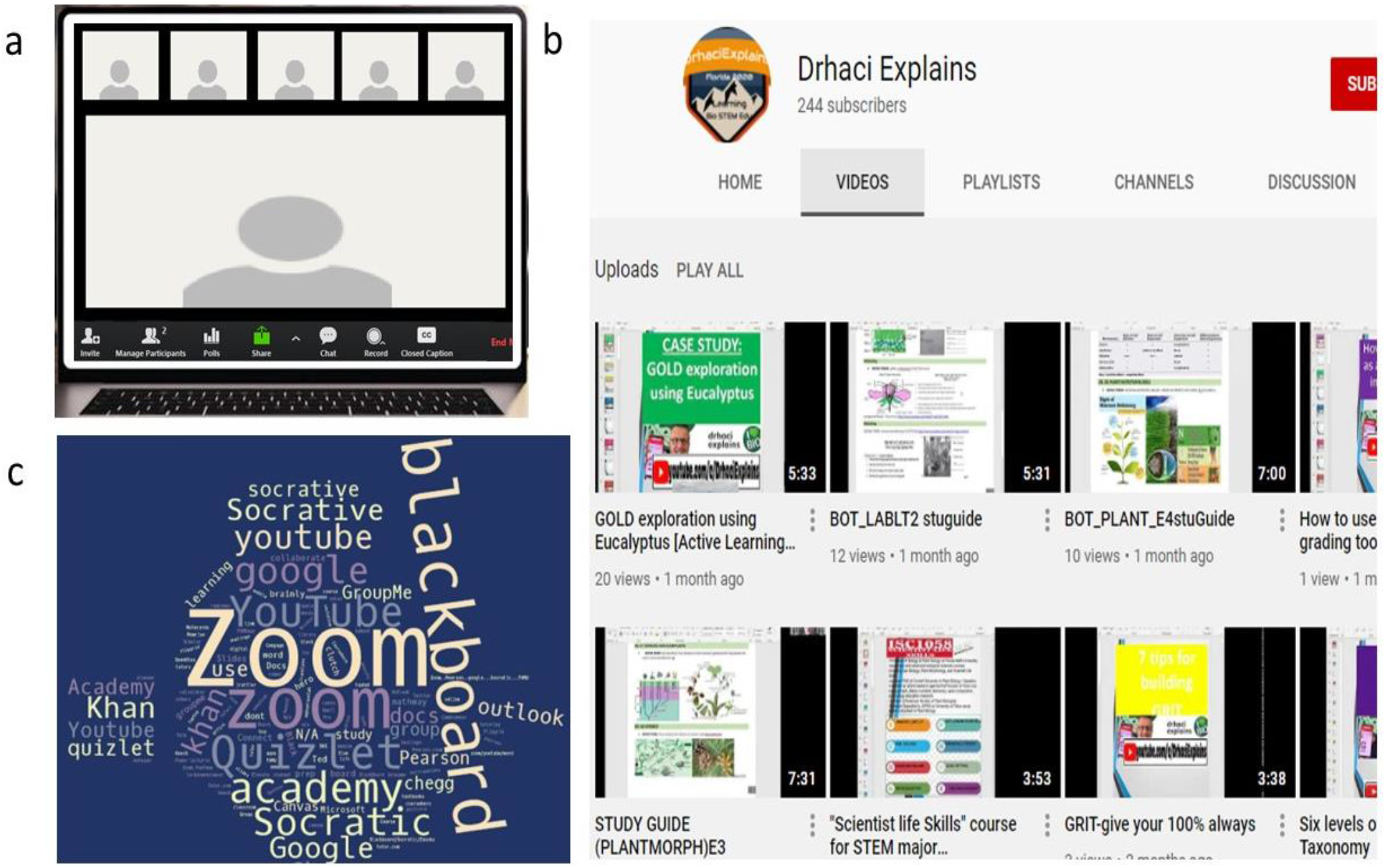
(a) A synchronously online class meets in a ZOOM room; (b) A YouTube channel developed in the Spring 2020 semester (https://www.youtube.com/c/DrhaciExplains) dedicated to STEM courses; and (c) A word-cloud of the total keywords cited by Spring 2020 cohorts in the survey.

### Survey Process, Participants, and Outcome Measures

The survey of student experiences in this new normal time and COVID-19’s impact on learning was designed as an electronic survey (Tables 1 and 2). The survey consisted of 10 statements and 11 questions. The answer options for the statements ranged from 1 to 5 (1= Strongly Disagree, 2= Disagree, 3= Neutral, 4= Agree, and 5= Strongly Agree). The remaining survey questions were open ended and multiple choice (MC) types. All survey items were reviewed by researchers with undergraduate teaching and assessment expertise. The response rate was approximately 88% of the students enrolled.

**Table 1.** Summary of the Descriptive Information of Students Enrolled in BSC1010 and BOT3303 in Spring 2020

| <u>Gender</u> |  |
| --- | --- |
| Female | 74% |
| Male | 26% |
| <u>Other student attributes</u> |  |
| First generation college student | 33% |
| Receive A's | 45% |
| Attend all classes | 65% |
| Enjoy learning | 74% |
| Work hard | 77% |
| Have access to reliable internet | 82% |

**Table 2.**
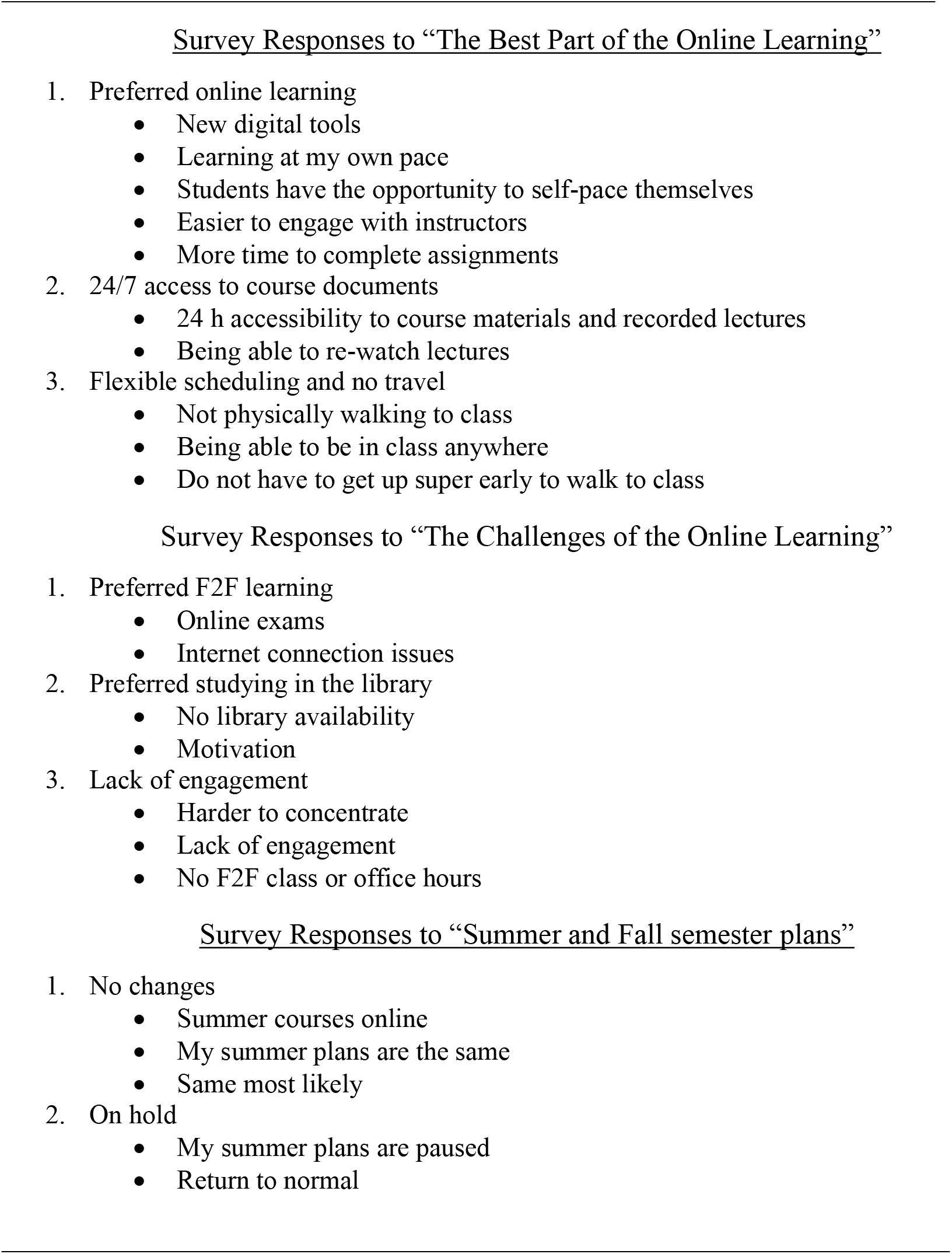
Survey Results of the Perceptions of the Students Enrolled in the Spring 2020

Class attendance was recorded both before and after pandemic for the F2F and online modalities, respectively. The two courses surveyed in this study were the following: General Biology I (BSC1010) and Plant Morphology (BOT3303).

### Ethics Statement

The Institutional Review Board at the university reviewed and approved this study under “IRB Approval Number 1597667-1”.

## Results

### Instruction Modality

Face-to-face (F2F) instruction is a live teaching modality in which the students and instructor are physically present in a classroom space (Fig. 2). On the other hand, online instruction is a virtual teaching modality via the internet that could be synchronous or asynchronous with a more flexible scheduling (Fig. 2). For centuries, universities have been using the F2F instruction modality. In the Spring 2020 semester, our courses shifted to online in mid-March. Therefore, Spring 2020 has emerged as an unprecedented dual modality semester with F2F and online.

### General Characteristics and Demographics of Courses

Table 1 summarizes the demographic information of those surveyed in this study. Overall, the students were biology, premedicine, or pharmacy majors. The survey included data from the General Biology I (BSC1010) and Plant Morphology (BOT3303) courses during the Spring 2020 semester. The majority of the participants were female (74%). While 33% of the participants were first-generation college students, 65% attended all classes (Table 1). Based on the Spring 2020 semester, the average class attendance increase was 9% in BSC1010, but there was not a significant increase in BOT3303 (Fig. 4). Moreover, the majority of the respondents stated that they enjoyed learning (Table 1).

**Figure 4.**
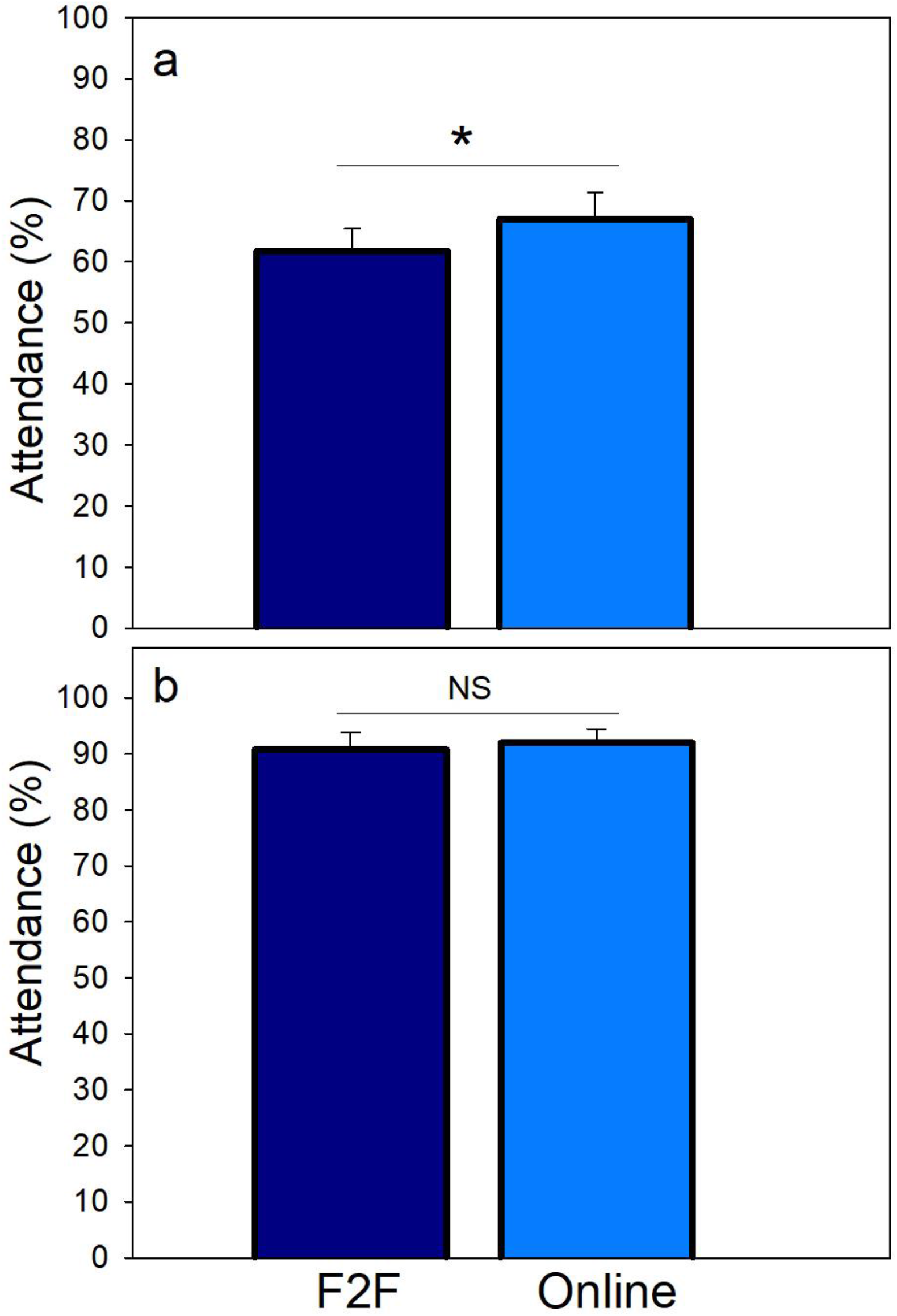
Displays of the student attendance of the two modalities. (a) General Biology I (BSC1010); and (b) Plant Morphology (BOT3303). F2F: face-to-face modality. Means were significantly different (Student’s t-test, p < 0.05) for (a) but not significantly different for (b).

### Student Perceptions of Online Learning

The Florida higher education board directed all universities to move classes online to slow the spread of the coronavirus in mid-March. In general, the majority of the participants (88%) reported that the current COVID-19 pandemic climate impacted their education (Fig. 4). The survey revealed that 79% of the participants reported that the F2F in classroom instruction format was the best modality for them. Students stated that they have taken online courses before this semester and they like having 24/7 accessibility to the course material in their own environment as online learners (Fig. 5).

**Figure 5.**
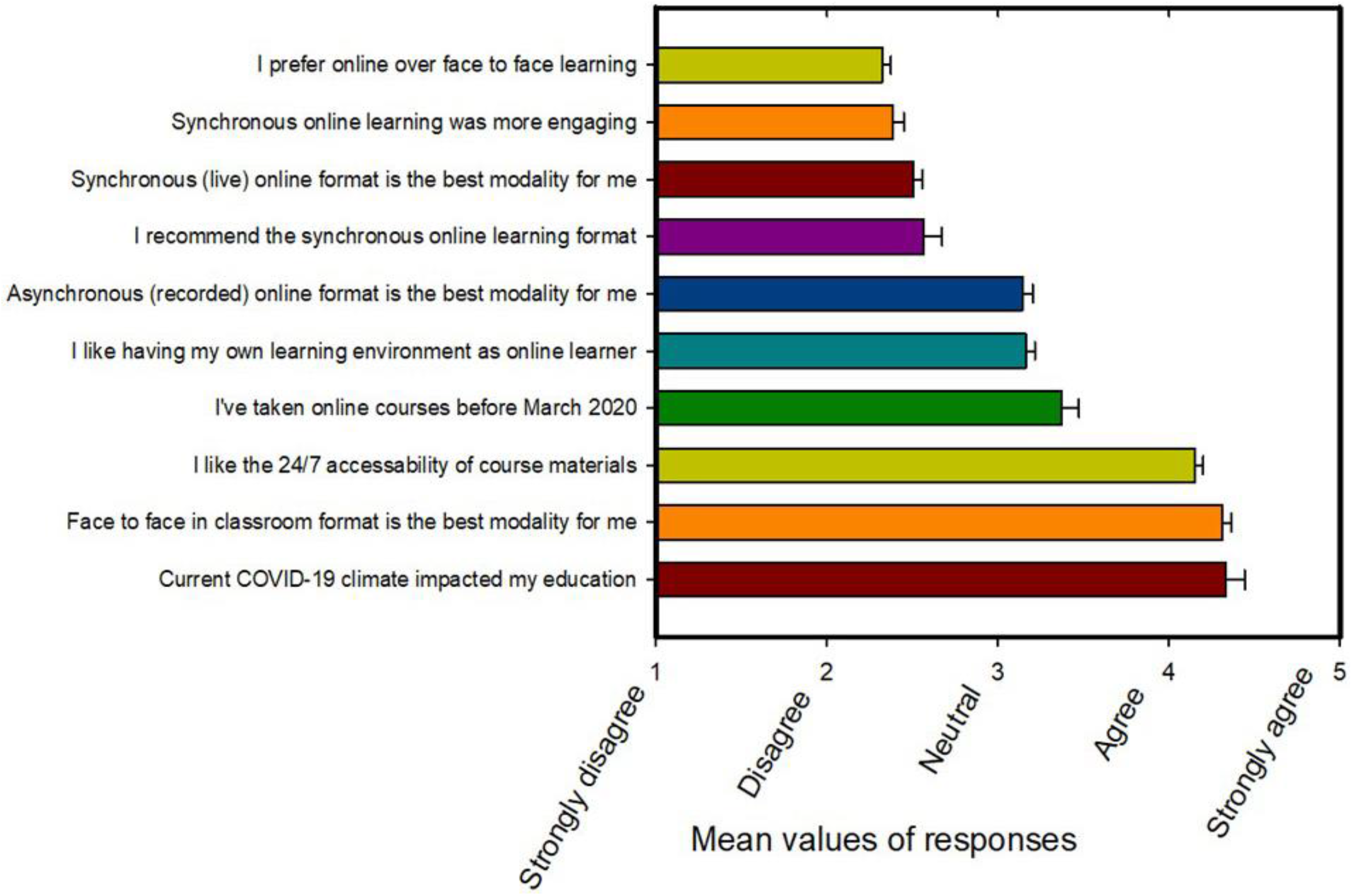
Student COVID-19 pandemic experiences survey and mean scores of each survey item with standard errors (SEs).

Our survey also included the perceptions of students which are summarized in Table 2 under four themes. In response to a question on the best part of online learning, students enjoyed 24 h accessibility to course documents, recorded lectures, and self-paced learning. As a response to the challenges of online learning, students reported motivation, concentration difficulty, and internet connection issues. It should be noted that student participants did not report major changes in their upcoming Summer or Fall semester plans (Table 2).

## Discussion

Extraordinary unprecedented circumstances are being endured during the deadly coronavirus pandemic (COVID-19). As a result, universities adapted to the new era by moving teaching online. Video conferencing technologies such as Zoom (ZM, San Jose, CA) allowed the live delivery of lectures and laboratories remotely.

In a challenging COVID-19 pandemic environment, both F2F and online instruction were taught by faculty and taken by the same students in the same semester (Spring 2020). In terms of class attendance, our study showed a modest increase of 9% in General Biology I (BSC1010) but no significant differences in Plant Morphology (BOT3303) (Fig. 4). Although it is difficult to interpret the attendance pattern differences, it is assumed that no travel time due to the pandemic and less distraction should result in improved remote attendance. Moreover, freshman class students were more likely to participate in online classes compared to upper class students in Plant Morphology (BOT3303).

The online modality allowed the instructor to connect distantly, broadcast both classes (BSC1010 and BOT3303) synchronously to all students via Zoom (Fig. 3a)), and record short videos (~6 min duration) to share on the newly developed STEM channel (DrhaciExplains, 2020; Fig. 3b). Furthermore, a majority of students also agreed that they have utilized digital applications such as Zoom, Blackboard, Socrative, YouTube, and Khan Academy among others (Fig. 3c).

Student comments concentrated on four major themes. Our survey results demonstrated that while the majority of students prefer the F2F instruction and learning approach, they also liked having access in their own environment, the 24/7 accessibility of course materials, and videos (Fig. 5). This is consistent with the findings of previous studies in Australia, which showed that both modalities delivered the same overall benefits to students (Ifediora, 2019).

When asked what they liked the least about online learning, student comments highlighted their lack of concentration, internet connection issues, and the lack of library availability (Table 2). When asked about future plans, student comments did not point to any major changes (Table 2). These results mirror previous studies where student attitudes regarding online and F2F did not differ at the University of Wisconsin (Nennig et al., 2020).

As the ongoing COVID-19 pandemic’s spread was detected statewide, in accordance with the guidance issued by the State University System, the university moved classes online by March 13 (Fig. 1b). The results presented in this study are promising and contained data from two biology STEM courses at a public HBCU in Florida, USA. Therefore, the Spring 2020 semester represented two equally relevant sections of F2F and online learning.

## Conclusions

This study investigated the impacts of COVID-19 on the undergraduate education, therefore it enhances the literature regarding F2F and online learning experiences. In conclusion, the current research fills the gap in the existing literature by providing the first study on the effects of the ongoing COVID-19 pandemic on undergraduate learning and experiences in the most unique dual modality of the Spring 2020 semester. Further research is recommended to examine what is occurring in other educational settings due to the impact of COVID-19 pandemic.

## Acknowledgements

The author gratefully acknowledges certificate of the excellent technical editing and reviewing of this manuscript provided by “American Chemical Society Author Services”. Special thanks are given to all enrolled undergraduates in the Spring 2020 in this unprecedented time finishing their studies when the world has changed. The authors also thank to Office of Instructional Technology for their help with online content design and virtual teaching tools. The author is grateful to the journal editor and reviewers for their valuable feedback during the review process.

